# Single-Particle Tracking and Positional Phenotyping Reveals Variant-Specific Early Checkpoints in SARS-CoV-2 Cell Entry

**DOI:** 10.64898/2025.12.18.695048

**Authors:** Frank H. Schulz, Marcus W. Dreisler, Denis Koylyu, Julián Valero, Mette Galsgaard Malle, Laia Civit, Jørgen Kjems, Nikos S. Hatzakis

## Abstract

SARS-CoV-2 entry is governed by Spike (S) protein-mediated engagement of ACE2 and subsequent activation of either plasma membrane fusion mediated by TMPRSS2 or endocytic uptake. Currently, most insights into these pathways come from bulk assays that obscure the fate of individual virions, thereby concealing intricate mechanistic details that can inform on therapeutic intervention strategies. Here, we applied single-particle fluorescence imaging to directly observe the early checkpoints of SARS-CoV-2 cell entry pathways and separate binding from internalization. Fluorescent virus-like particles (VLPs) pseudotyped with either G614 or Omicron BA.5 S protein variants were imaged on HEK293T-ACE2 (TMPRSS2-negative) and classified at the single-particle level as surface-interacting, crossing, or internal. At baseline, G614 VLPs show higher binding and a larger internalized share than BA.5 VLPs, revealing general divergence in early entry behavior between variants. A trivalent anti-S receptor-binding domain aptamer reduces G614 binding and lowers its internalization. Conversely, the aptamer does not block BA.5 VLP cell binding but increases its internalization efficiency. Pitstop 2, an inhibitor of clathrin-mediated endocytosis, causes no significant change in this observation window, consistent with early clathrin-sensitive events having already progressed. Quantification of trajectories reveals variant-specific mobility: BA.5 displays higher step length than G614, consistent with greater lateral scanning and surface retention. Together, these compact single-particle readouts expose variant-resolved early checkpoints in entry and provide a simple platform to test how ligands and pathway probes shift binding and internalization.

## INTRODUCTION

SARS-CoV-2 is an enveloped betacoronavirus whose trimeric Spike (S) protein mediates host-cell entry by binding ACE2 and undergoing proteolytic activation by shedding of its S1 component. Cellular entry can proceed at the plasma membrane when the protease TMPRSS2 is available or after endocytosis when activation is supplied by endosomal cathepsins, and balance is cell-type dependent (1). With Omicron variants, several groups showed a shift away from a TMPRSS2-dependent entry mechanism toward endosomal, cathepsin-dependent activation, reshaping tropism and fusion location (2,3). In TMPRSS2-negative systems such as HEK293T-ACE2, drug-based perturbations and gene-based manipulations consistently indicate acidification-dependent endosomal entry (4–6).

Mutations in the S protein have been shown to strongly influence viral entry mechanism. The D614G change (G614) stabilizes the trimer in its open receptor-binding state, reduces S1 shedding, and increases functional S protein density on particles, boosting entry without necessarily raising monomeric ACE2 affinity (7,8). Within Omicron, BA.5 maintains robust ACE2 engagement and altered protease preferences relative to pre-Omicron variants, with context-dependent consequences for cell entry and fusogenicity (2,9).

Coronaviruses are known to exploit multiple internalization routes. Clathrin-mediated endocytosis (CME) has been implicated in SARS-CoV-2 internalization (10). Yet clathrin-independent pathways and cytoskeleton-guided delivery are also observed, including filopodia-assisted transport to endocytic hot spots (11) and direct co-localization with clathrin-coated pits alongside diverse pre-internalization motion states (12). Deciphering cell entry pathways is crucial for rational design of entry-targeting antivirals as they will only be effective if they target the actual pathway(s) used by the specific virus variant. This is especially challenging in live cells, where overlapping pathways, fast-acting mechanisms over small length scales obscure overlapping entry pathways.

To directly observe how individual viruses bind the cell surface and progress towards cell entry, a range of single-particle imaging approached have been used. Single-particle methods solve many of the limitations of bulk assays by providing high spatial and temporal resolution on individual viruses. Total internal reflection (TIRF)-based single-particle tracking (SPT) can dissect search, capture, and confinement at the surface (13), but it cannot follow particles once they move out of the evanescent field during internalization, leaving the actual transition into endocytosis unresolved. Active-feedback 3D tracking follows freely diffusing virions as they make first contact and transition into the cell (14) but provides limited quantitative insights as only a single virus particle can be tracked at a time. Label-free and correlative approaches further map motion states of the endocytic structures (12), yet they do not provide the number of bound particles that proceed to internalization. Quantifying this internalization fraction, the proportion of surface-bound viruses that internalize within a defined time window, and how it depends on virial variant or pathway perturbation, is essential for mapping the full cell entry pathway and for rational design of entry-antivirals.

In this study, we built a toolkit to quantify cell entry and its dependence on mutation and potential inhibitors that perturb entry. We employed bright fluorescent, non-infectious virus virus-like particles (VLPs) built on the HIV Gag-EGFP scaffold, which present S protein in its native conformation and enable safe, quantitative assays (15). We combined these with a trivalent anti-receptor binding domain (RBD) aptamer as a high-affinity, multivalent modulator of S protein-ACE2 engagement (16), and with Pitstop 2 as a probe of CME (17). Using these tools together, we performed variant-resolved, single-particle measurements to directly quantify binding, surface mobility, and internalization during the early moments of SARS-CoV-2 entry. Besides providing mechanistic insights on viral entry, the method can be used as a screening platform for antiviral compounds.

## RESULTS

### A fast positional assay defines early entry

We developed and implemented a single-cell-based single-particle assay designed to isolate the internalization step of entry. Using HEK293 cells overexpressing ACE2 (HEK293T-ACE2) but lacking TMPRSS2, internalization was restricted to the endosomal route (Fig. 1a). We classified individual VLPs as surface VLPs, crossing VLPs, or internal VLPs to capture distinct stages of early entry (Fig. 1a). This simple spatial framework separates cell surface binding from internalization, allowing quantitative comparison of how variants or perturbations shift binding and internalization dynamic. The fractional distribution of particles among these positions directly reports how many virions remain bound, cross the cell membrane, or accumulate internally, providing an intuitive and quantitative readout of early entry behavior without complex trajectory modeling. Here, we focus on the fraction of all surface-bound VLPs that proceed to internalization during the movie, referred to as the internalization fraction.

**Fig. 1:**
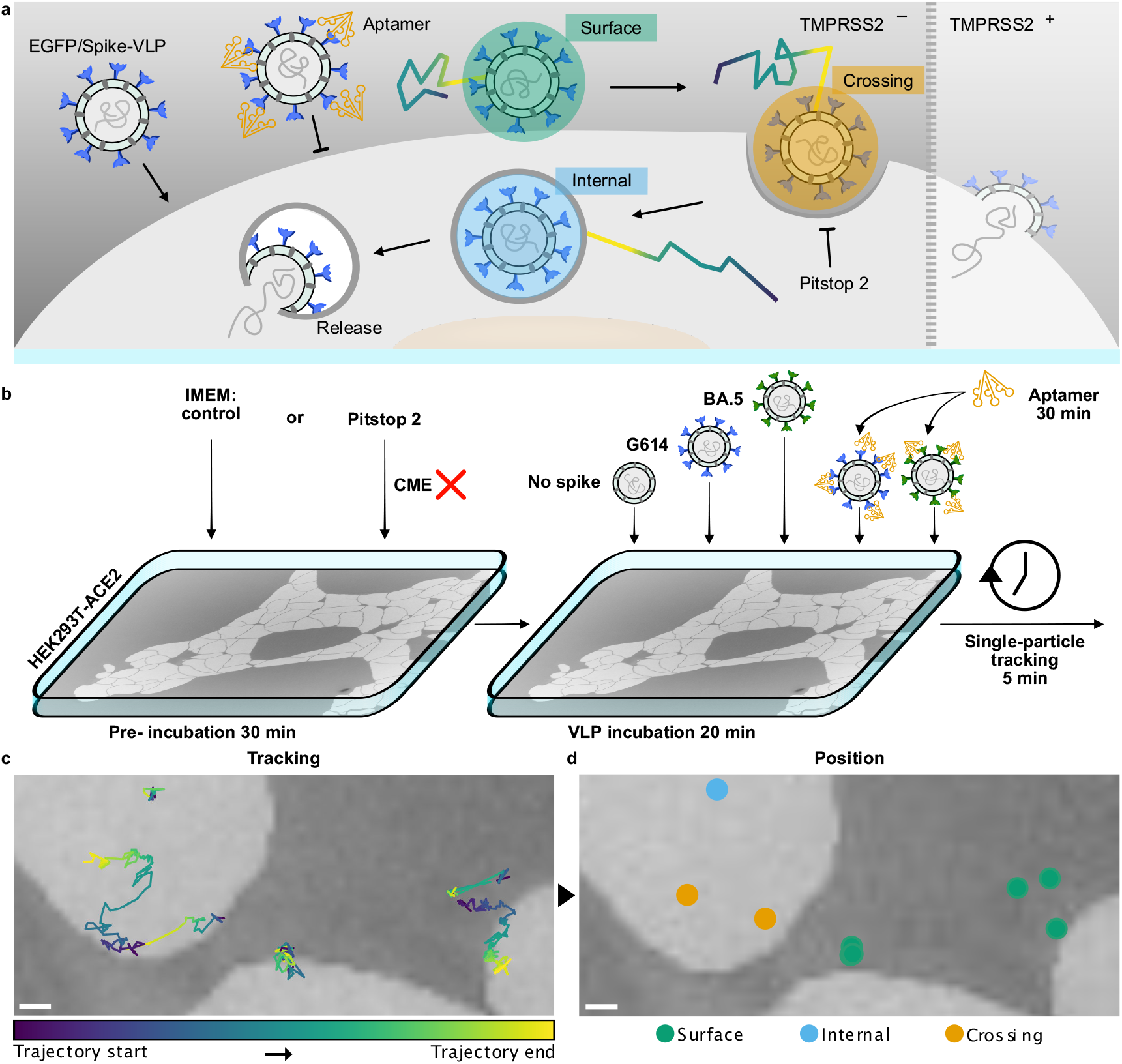
Schematic representations of experimental assay, and analysis to study SARS-CoV-2 cell entry. **a**, Basis of SARS-CoV-2 cell binding and entry, and use aptamer and Pitstop 2 for probing binding and entry. Enveloped VLPs assembled using HIV Gag-EGFP fusion protein and decorated with SARS-CoV-2 S proteins bind to the surface of cells. TMPRSS2-negative HEK293T-ACE2 were used to prevent direct fusion with the outer cell membrane. A trivalent anti-S aptamer can selectively bind to the VLPs and inhibit their binding to cells. VLPs that bind to the cell surface (green mark) can be immobilized in one spot or scan the surface for entry sites. For cell entry, VLPs must cross the cell membrane through endocytic pathways (orange mark). Fully internalized VLPs (blue mark) will stay in endosomes until fusion with the endosomal membrane where native viruses would release their viral RNA (dark grey strands). **b**, Overview of experimental conditions. HEK293T-ACE2 cells were incubated in either control medium (IMEM) or in IMEM with Pitstop 2 to block endocytosis for 30 minutes before addition of VLPs. To IMEM cell, one of each type of VLP was then added: no S protein control, G614, BA.5, G614 preincubated with trivalent aptamer for 30 minutes, or BA.5 preincubated with aptamer for 30 minutes. To Pitstop 2-treated cells, either G614 or BA.5 VLPs were added. Under all conditions, cells were incubated for 20 minutes with VLPs for recording of SPT videos for 5 minutes. **c**, Representative cell assay area of VLP trajectories from recorded data overlayed masked cell areas (light grey) and non-cell areas (dark grey). Trajectories are color coded by their individual time progression from start of trajectory to end of trajectory. **d**, Positional phenotyping of VLPs. Based on spatial positions relative the masked cell membrane, VLP trajectories from **c** are individually classified as either one of three positional identities: *surface* which only reside on the cell membrane; *crossing* which convert from being membrane bound to internalizing; or *internal* which only reside inside the cell lumen. Scale bars, 1 µm.

To ensure that our measurements reflect events that happen between cell binding of VLPs and early states of endocytosis, all measurements were made within 25 minutes after the initial addition of VLPs to cells (Fig. 1b). This timeframe includes a 20-minute incubation to allow VLPs in solution to bind to the cell surface and for internalization to initiate, followed by 5 minutes of active recording and single-particle tracking using spinning-disk confocal microscopy. This captures a defined window of viral entry events before advanced endosomal maturation and eventual fusion-mediated release (4–6). In this TMPRSS2-negative context, the “internal” state corresponds to a pre-fusion endosomal pool, so the internalization fraction is an early entry readout rather than a measure of downstream fusion or replication (4).

We extracted the spatial localization of individual diffraction limited VLPs (18,19) and extracted their single-particle trajectories (Fig. 1c) that quantitatively describe their motion. As argued previously (20,21) motion can encode both identity and spatial localization. For each step in a trajectory, we determined whether the VLP was inside or outside the cell and computed its distance to the nearest segmented cell membrane (Supplementary Fig. 1). Using a motion-informed, distance-based threshold, we assigned each trajectory to a single VLP and a single positional identity (Fig. 1d; Supplementary Fig. 2). This classification allows for direct comparison of VLP counts across positions between conditions and enables analysis of diffusional behavior at each position.

We evaluated how viral cell entry pathways are affected by two compounds with different targets along the entry pathway and different variant specificities. A trivalent anti-S protein aptamer targeting RBD (Fig. 1a,b) (16) was used to study VLP cell binding and the dependence of RBD availability on VLP internalization. This aptamer was reported to bind early SARS-CoV-2 variants with proven binding selectively to and neutralization of G614 S protein (16) while the monomeric version of the aptamer showed near background binding to Omicron BA.2 S protein which is a close relative of BA.5 S protein (22). For these reasons, the trimeric aptamer can be used to probe variant-specific effects on binding and internalization. Pitstop 2 is an inhibitor of CME (17) and was used to test pathway dependence (Fig. 1a,b).

### G614 shows higher binding and internalization than BA.5 at 20 min

We first compared total binding and early entry behavior between VLP variants under unperturbed conditions in imaging medium without inhibitors (IMEM) by counting the total number of VLP trajectories at all three classified positional states. Because this number is a convolution of bound and internalized VLPs, it reflects all sustained binding events that have taken place in the 25-minute experimental timeframe (Fig. 2a), where each dot represents a single VLP trajectory and colors indicate its positional identity: green for surface, orange for crossing, and blue for internal.

**Fig. 2:**
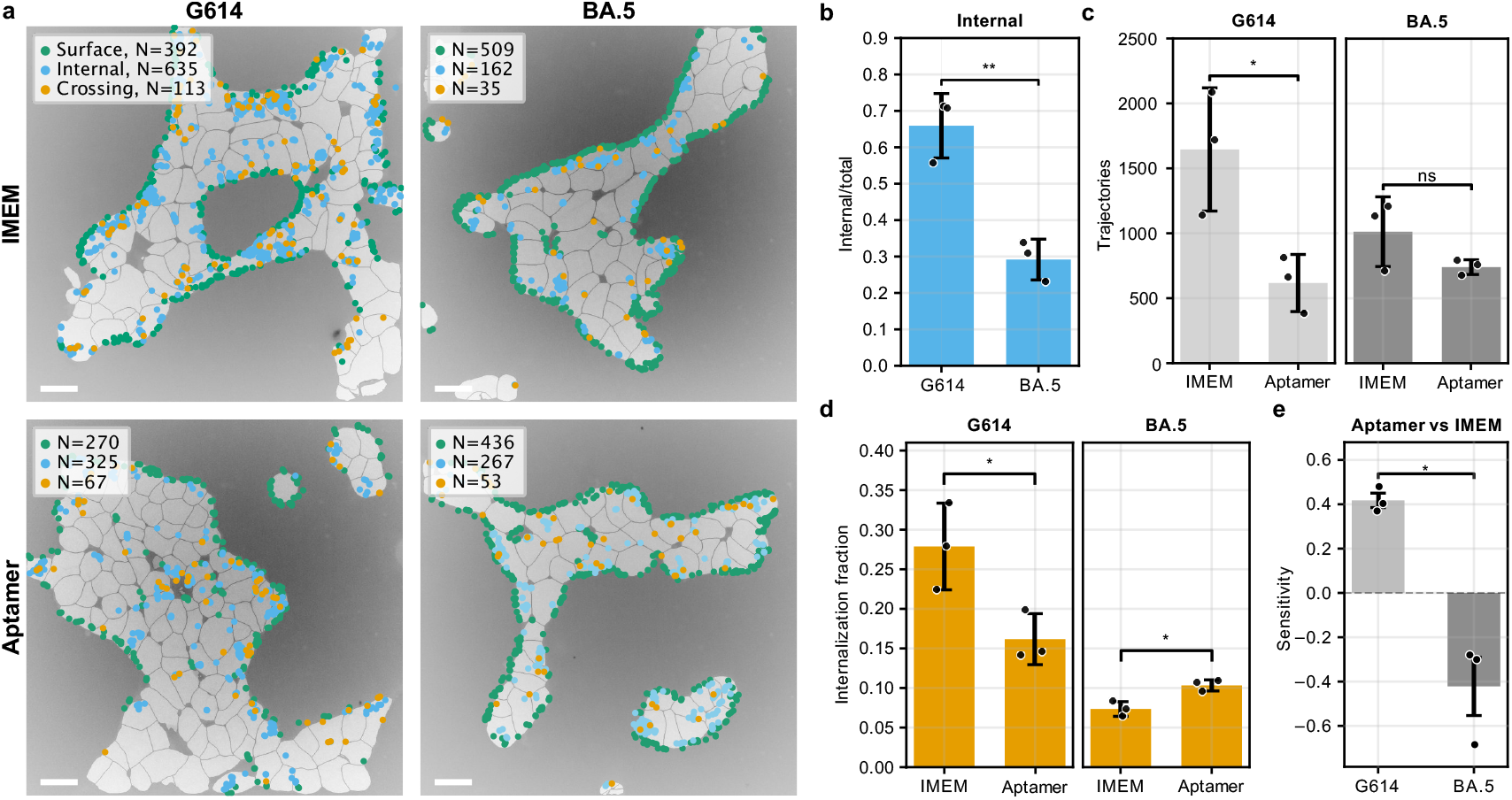
Effect of trivalent anti-S aptamer on VLP binding and entry. **a**, Positional phenotyping data G164 and BA.5 VLPs in cells with or without trivalent aptamer. Individual dots represent single VLP trajectories and are color coded according to their positional identity. One of three biological replicates is shown per condition. **b**, Fraction of internalized G614 and BA.5 VLPs in IMEM. **c**, Quantification of total number of VLPs (sum of surface, crossing and internal) of conditions represented in **a**. and thus reflect all binding events throughout the total 25 minutes the cells are exposed to VLPs. **d**, Internalization fraction of G614 and BA.5 VLPs treated or not treated with aptamer, where the internalization fraction denotes the fraction of surface VLPs converting to internal. **e**, Aptamer-sensitivity of internalization of VLPs treated or not treated with aptamer. The index is based on the internalization fraction in **d** and describes how the internalization of VLP is affected by aptamer relative to no aptamer addition. Positive values denoting an inhibitory effect on internalization and negative values denoting an enhancing effect on internalization. Trajectory count bars show mean ± SEM of three biological replicates while all other bars show mean values calculated from sample means and SEM with propagated errors (*N*=3). Scale bars, 20 µm. Significance, ns = p>0.05, * = p<0.05.

Between conditions (different S protein variants and inhibitors), VLPs were added to cells in equal concentrations, ensured by VLP protein quantification and NTA measurements. First, we see that binding of VLPs to HEK293-ACE2 cells is S protein-dependent as naked VLPs show almost no trajectory counts on cells relative to S protein-decorated VLPs (Supplementary Fig. 3). Second, despite a lower total binding of BA.5 VLPs compared to G614 is observed, the difference is not significant (Supplementary Fig. 3).

The fractional distributions of VLPs at the three cellular positional states show highly different tendencies between G614 and BA.5 with nearly opposite distributions of surface and crossing fractions (Fig. 2b and Supplementary Fig. 3). Here, G614 VLPs have a much higher fraction of internalized VLPs than BA.5 reflecting a higher degree of internalization and retention in pre-fusion vesicles (Fig. 2a,b). Likewise, BA.5 VLPs show much higher surface retention than does G614 in the observed timeframe. This may seem contradictory since BA.5 is more endocytosis-dependent than G614. However, it is important to note that the size of the internal fraction is not a measure of cathepsin-mediated fusion and release from endosomes for infection.

### Anti-S protein aptamer alters VLP binding and internalization in a variant-specific manner

To investigate how blocking of the S protein RBD affected VLP binding and internalization, we used a trimeric anti-S aptamer which was selected against Wuhan strain S protein (Fig. 2a) (16,22). This variant specificity is clearly reflected in the total trajectory count, where G614 counts are reduced by 2/3 while there is no significant change in the trajectory count for BA.5 (Fig. 2c).

We then examined the internalization fraction, defined as 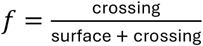, which decreases for G614 but interestingly increases for BA.5 with aptamer functionalization (Fig. 2d). We summarize that with the aptamer-sensitive fraction on internalization, 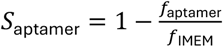, which is positive for G614 and negative for BA.5 (Fig. 2e). Together, these results show that the aptamer suppresses internalization during early entry in G614 but enhances it in BA.5, revealing opposing effects of the same ligand on early uptake across variants (Fig. 2a,c–e). VLP intensity distributions are narrow without the appearance of bright tails, arguing against clustering and avidity-induced internalization as the explanation (Supplementary Fig. 4).

### Pitstop 2 does not significantly change in internalization or CME index

To test how clathrin contributes to internalization in our setup, we added Pitstop 2 and evaluated internalization in the same positional framework (Fig. 3a) (10,17). Under Pitstop 2, the internalization fraction shows no significant change for either variant (Fig. 3b). We summarize the perturbation with the CME-sensitive fraction 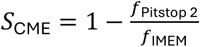: values hover near zero, with a small positive tendency for G614 and a small negative tendency for BA.5, neither significant in our dataset (Fig. 3c). Since our imaging happens after a 20-min pre-incubation, these are conservative, late-window estimates of clathrin involvement: any strongly clathrin-dependent events that occurred early are likely already counted in the internal VLP pool by the time we start imaging (4). Given reported off-target effects of Pitstop 2 beyond clathrin terminal-domain engagement, we interpret these late-window CME-sensitivity estimates conservatively (23,24).

**Fig. 3:**
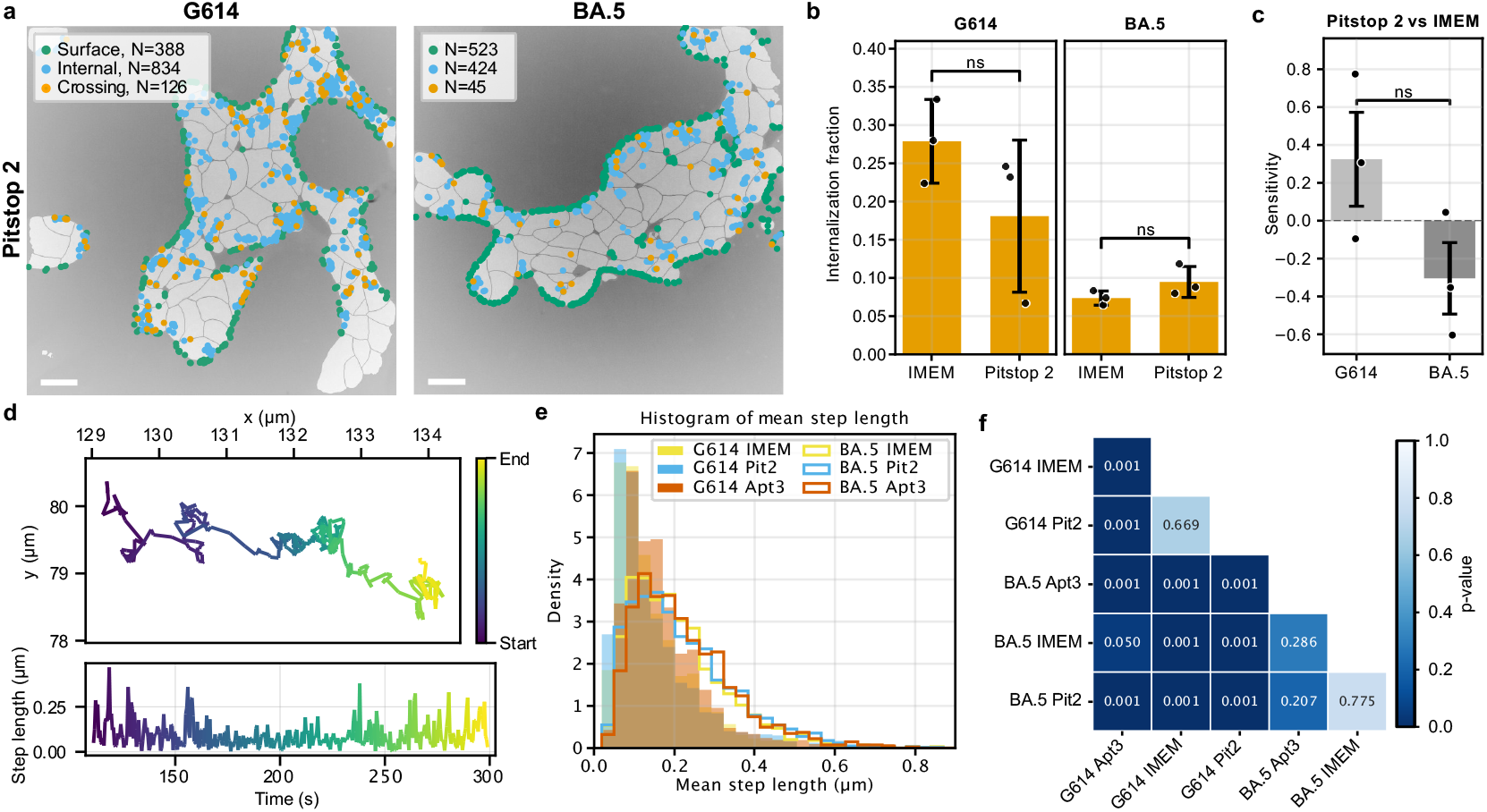
VLP sensitivity towards Pitstop 2 and pre-internalization diffusion. **a**, Positional phenotyping data G164 and BA.5 VLPs in cells treated with Pitstop 2. **b**, Internalization fraction of G614 and BA.5 VLPs in cells treated or not treated with Pitstop 2. **c**, CME-sensitivity of internalization VLPs in cells treated or not treated with Pitstop 2. The index is based on the internalization fraction in **b** and describes how the internalization of VLP is affected by Pitstop 2 relative to control. Positive values denoting an inhibitory effect on internalization and negative values denoting an enhancing effect on internalization. **d**, Example of VLP trajectory used to investigate differences in diffusional behavior in different experimental conditions. **d top**, Each trajectory is described by x- and y-coordinates per frame connected by a step. **d bottom**, Kymograph of step lengths of the trajectory shown above. Higher step lengths imply faster movement of the VLP between two points. Trajectory segments with high motility show high step lengths and vice versa. **e**, Histogram of step-length distributions of G614 (filled) and BA.5 (steps) surface-classified VLPs under all experimental conditions. **f**, Heatmap of pairwise p-values for the comparisons in **e**. p-values come from a likelihood-ratio test under a log-normal model with a parametric bootstrap, computed on balanced subsamples (n = 200 per group) to avoid large-sample inflation. *N*_G614,IMEM_ = 4936, *N*_G614,Pit2_ = 3674, *N*_G614,Apt3_ = 1851, *N*_BA.5,IMEM_ = 3039, *N*_BA.5,Pit2_ = 3878, *N*_BA.5,Ap3_ = 2220. Low p-value indicates different surface step-length distributions; high p-value indicates no detectable difference at this sample size. Bars show mean values calculated from sample means and SEM with propagated errors (*N*=3). Scale bars, 20 µm. Significance, ns = p > 0.05, * = p < 0.05.

### Surface mobility is variant-specific: BA.5 VLPs show higher surface step length than G614

To check whether our perturbations changed VLP motion, we examined step length along single VLP trajectories and then the population summaries. The example trajectory alternates between fast and slow segments. High step lengths mark rapid lateral movement (18,19,22,23), low values mark confinement (Fig. 3d) (27,28). Here, we found that G614 and BA.5 exhibit different behavior at the cell surface but not in their crossing or internal fractions (Supplementary Fig. 5).

Pooling surface segments, the mean step length distributions are most similar within each variant and separated between variants (Fig. 3e). All BA.5 conditions overlay closely and sit to the right of G614, consistent with faster surface motion for BA.5. G614 is likewise stable across IMEM and Pitstop 2, with a modest shift only in the presence of aptamer (Fig. 3e).

We formalized these observations with a likelihood-ratio test on balanced subsamples. Between-variant comparisons at matched conditions were significant, confirming the consistent BA.5 shift to higher step lengths, whereas within-variant comparisons were largely not significant: high p-values for BA.5 across treatments and for G614 between IMEM and Pitstop 2, with a lower value only under aptamer treatment (Fig. 3f). This selective G614 response aligns with its stronger aptamer engagement, indicating that surface mobility reflects variant-specific behavior rather than drug effects (Fig. 2a,c–e; Fig. 3d–f).

## DISCUSSION

In summary, we leveraged a complementary toolkit to quantify the internalization fraction of VLPs and delineate its dependence on both viral variants and entry inhibitory compounds at the single-particle and single-cell level. Our data collectively support that the G614 variant shows a larger bound population and a larger internal share than the BA.5 variant within the same observation window. Prior work links G614 to reduced S1 shedding and greater S protein trimer stability (7), which could plausibly increase the probability that bound particles proceed to internalization. In TMPRSS2-negative cells like HEK293T-ACE2, activation is expected to occur in acidification-dependent endosomes rather than at the surface, so a larger internal share reads as internalization and retention in a pre-fusion endosomal pool within our window (1,4–6).

The trivalent anti-S protein RBD aptamer separates early entry checkpoints by S protein variant. For G614, strong aptamer interaction with the S protein reduces VLP cell binding and lowers internalization within the observation window. Conversely, for BA.5, binding remains essentially unchanged while internalization increases. The narrowing of VLP fluorescence intensity distributions upon aptamer addition argues against clustering and avidity effects as drivers of uptake. These opposite responses align with variant-specific aptamer affinity: effective RBD blockade on G614 limits both particle accumulation and uptake, while weaker RBD recognition on BA.5 leaves most spikes accessible and may permit additional electrostatic or RNA-mediated contacts reported for Omicron S proteins (29,30), favoring recruitment through alternative, non-ACE2 routes (2,3,16,31–33).

Surface mobility of VLP adds a layer of mechanistic understanding to the counts and positional distributions. Across conditions, BA.5 exhibits higher surface step length than G614. Faster lateral motion is consistent with extended surface scanning before internalization, which can increase the BA.5 surface pool within the fixed window while leaving total binding comparable. Within variant, mobility is stable between IMEM and Pitstop 2 as only G614 shifts under aptamer, which is exactly where binding specificity predicts an effect.

Pitstop 2 produces little, and non-significant, change in internalization in our hands. Two features of the assay make this a conservative, late window estimate rather than a negation of clathrin. There are several hypotheses that my explain our observations: first, imaging begins after a 20-minute pre-incubation, so early, strongly clathrin-dependent entries may already have occurred; second, we do not wash after addition of VLPs, so new VLPs can still bind during imaging. Because Pitstop 2 is present continuously in the VLP medium, inhibitor concentration is constant, but the ongoing binding of VLPs can enrich the surface pool with late binders that have had less time to internalize in the experimental timeframe. Pitstop 2 targets the clathrin terminal domain but has reported off-targets and can influence clathrin-independent internalization, which tempers interpretation (17,23,24). Prior studies implicate CME in SARS-CoV-2 internalization across systems (10). Our data instead reveal that during the late entry window captured here, internalization proceeds largely through clathrin-independent or mixed pathways, underscoring the method’s sensitivity to pathway timing.

The presented approach is an efficient single-molecule platform to screen antiviral compounds and simultaneously study their mechanism of action and the extracted information is directly applicable for entry-stage intervention. Because binding and internalization are separated, probes and antiviral therapeutics can be classified by dominant action within minutes. Binding blockers (for example, soluble ACE2 or high-affinity RBD binders) should reduce total particle counts with modest effects on internalization among surviving particles as reflected by our G614 aptamer case (16,34). Internalization modulators that alter endosomal maturation or trafficking (for example, PIKfyve or endosomal pH pathway inhibitors) should primarily shift internalization at near-constant totals, providing a compact screen without overlapping with binding (35–37). The variant-dependent divergence under a single trivalent ligand also guides binder design: for Omicron-class S proteins, tuning valency/spacing/epitope coverage to both reduce binding and lower internalization, either by raising BA.5 RBD affinity or pairing RBD with a second epitope, would be a rational direction (16,38,39). Host-factor strategies also become relevant as co-receptors such as SR-B1 can augment early entry, Antagonizing such pathways should lower internalization at roughly constant binding in responsive cells, providing a signature the assay can detect (40).

The scope of the assay is intentionally focused on pre-fusion events. VLPs therefore report the early stages of entry rather than replication, with the internal fraction corresponding to a pre-fusion endosomal pool in this TMPRSS2-negative context, and internalization representing a defined kinetic transition within the observation window. State assignment is based on distance-to-membrane and positional persistence, which provides a conservative classification of crossing events while maintaining robustness across conditions. Although endocytic machinery was not co-labeled, the combination of timing and pharmacological perturbation already allows meaningful pathway discrimination, and future inclusion of tagged host factors could extend this framework toward direct pathway mapping (21,26).

Separating binding from internalization provides a universal metric for comparing viral variants, host factors, and therapeutic interventions across experimental systems. Since this method is widely applicable to any particle that must bind, internalize, and traffic within cells, these positional and kinetic readouts create a shared framework for analyzing viral entry, endocytic routing, and nanoparticle delivery. By turning nanoscale motion into quantitative entry outcomes, the approach extends beyond virology to illuminate general principles of how biological and synthetic particles interact with cells.

## Limitations of study

This study quantifies SARS-CoV-2 entry using a fluorescent reconstituted virus-like VLP system rather than authentic SARS-CoV-2 virions. S protein density, organization, and particle composition may therefore differ from infectious virus and could shift quantitative entry behavior. In addition, incorporation of HIV Gag to form VLPs may alter particle biophysical properties and intracellular trafficking relative to native SARS-CoV-2 particles.

Experiments were conducted in TMPRSS2-negative cells to emphasize endosomal uptake and focus measurements on pre-fusion events. Consequently, the entry mechanisms and inhibitor sensitivities observed here may not fully generalize to TMPRSS2-positive contexts such as airway epithelia, where plasma-membrane entry routes can contribute. Because the assay is intentionally restricted to pre-fusion readouts (binding, surface mobility, and internalization), it does not directly measure downstream fusion, uncoating, or productive infection.

Finally, VLP tracking was performed in 2D and positional phenotyping relied on distances to segmented cell boundaries. Out-of-plane motion, detection/tracking thresholds, and cell morphology/topography may therefore influence localization precision and derived metrics.

## METHODS

### Materials

All chemicals were of analytical grade and were purchased from Sigma-Aldrich Denmark, unless otherwise stated.

### Plasmids

GagEGFP - codes for HIV-1 GAG with EGFP fused to its C-terminal was a gift from Sven Eyckerman (Addgene plasmid #80605)(41). Spike D614G plasmid with 21 aa deletion in the C terminal was a gift from Jesse Bloom (42). Spike Omicron BA.5 was purhcased from SinoBiological.

### Production of pseudotyped fluorescent VLPs

VLPs were produced as described previously (15). In brief, VLPs were produced by calcium phosphate transfection of HEK293T cells with plasmids containing genomic sequences for HIV Gag-EGFP fusion and either or Omicron BA.5 variants of the SARS-CoV-2 S protein. VLPs were isolated by filtering cellular supernatant through a 0.45 μm filter and stored in the dark at 4 °C.

### VLP concentration measurement

High-binding half-area black microplates (Greiner Bio-One, cat. no. 675077) were used for total fluorescence measurements. A volume of 10 µL of each sample was dispensed per well, and fluorescence intensity was recorded at 479 nm excitation and 520 nm emission using a BioTek Synergy H1 microplate reader (Agilent).

### Aptamer preparation

Trimeric anti-S RBD aptamer was prepared as described previously (16). Prior to use, the aptamer was refolded to ensure correct structure in a thermocycler using a descending temperature ladder of 95°C for 2 minutes, 65°C for 5 minutes, 37°C for 2 minutes.

### Cell line

HEK293T-ACE2 (42) cells were maintained in growth medium consisting of DMEM high glucose supplemented with 1% penicillin-streptomycin and 10% FBS in 37°C and 5% CO_2_.

### Cell sample preparation

In preparation for SDCM experiments, cells were seeded at 15,000 cells per well in ibiTreat 8-well polymer microscopy slides (ibidi) in growth medium for 1 day. Biological replicates of experiments were performed on consecutive cell splits. Cell entry experiments started with a cell pre-incubation step by replacing growth medium with imaging medium (IMEM) consisting of DMEM high glucose without phenol red supplemented with 1% penicillin-streptomycin, 2% FBS, 2.5 mM L-Glu, and 3 mM MgCl2 for 30 minutes. For experiments involving Pitstop 2, IMEM was supplemented with 10 mM Pitstop 2 in the cell pre-incubation step. Both IMEM with and without Pitstop 2 contained 0.1% DMSO directly from or accounting for Pitstop 2 solubility. The pre-incubation medium was then replaced with identical medium containing equal concentration of one of the VLP variants. To IMEM cells, either naked VLP, G614 VLP or Omicron BA.5 VLP were added. Also, to IMEM cells, G614 VLP or Omicron BA.5 VLP that had been pre-incubated with 200 nM trimeric aptamer for 30 minutes at room temperature were added. To Pitstop 2 treated cells, either G614 or Omicron BA.5 variant S protein VLPs were added in IMEM containing 10 mM Pitstop 2. All VLP-containing medium also had 4 µM ATTO647-carboxy (ATTO-TEC) added for negative staining of cell outlines. All VLPs were added in equal concentrations. All cell experiments were conducted in biological triplicates on different days and different cell split numbers.

### Single-particle tracking of VLP in cells on SDCM

Microscopy was conducted on an Olympus SpinSR10 inverted SDCM equipped with a PRIME 95B CMOS camera (Teledyne Photometrics) and a 60× NA 1.4 oil immersion objective (Olympus), yielding a pixel size of 183.33 nm. ATTO647-carboxy contrast staining was excited with a 640 nm laser (10% power, 50.04 ms exposure), while EGFP-labeled VLPs were excited with a 488 nm laser (5% power, 50.04 ms exposure). SPT was performed for 5 minutes at a frame interval of 500 ms (600 frames total), with the 640 nm and 488 nm channels imaged consecutively for each frame.

### Imaging data

Two-channel time-lapse stacks were acquired, with one channel recording cell morphology *I*_*c*_(*x, y, t*), and the other capturing fluorescent particle signals *I*_*p*_(*x, y, t*). Particle positions (*x, y, t*) were detected in the particle channel for each frame, while cell channel was used for segmentation and subsequent distance analyses.

### Drift correction

Image stacks that displayed occasional field-of-view translocations (2 of 21 total movies) were corrected using the Fast4DReg (Legacy 2D) plugin in Fiji (43). Drift estimation was carried out on the fluorescence channel with time averaging disabled (set to 1), ensuring each frame was aligned independently and only abrupt shifts were corrected. The maximum expected displacement was set to 50 pixels, the first frame was used as reference, and corrections were applied directly to the dataset. Drift corrected files were cropped by 24 pixels on the x axis to remove empty space created during drift correction.

### Image processing and illumination correction

For each frame *I*_*c*_(·, *t*), a smooth background field 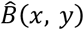 was estimated either by fitting a broad Gaussian surface or a polynomial surface. The corrected image was given by

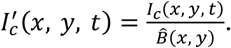

### Cell segmentation

Instance masks *M*(*x, y, t*) containing positive integer labels (1, 2, 3, …) for each distinct cell were generated using a pretrained Cellpose (44) model applied frame-by-frame. These were binarized as

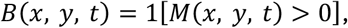

With outliers excluded to focus on analyses on cell interiors.

### Particle detection

SPT of VLP was performed with an in-house tracking script based on the Crocker-Grier algorithm using an estimated particle diameter of 9 pixels, ensuring also the tails of the point spread function were captured (45).

### Particle intensity extraction

The integrated particle intensity, *raw mass*, was background-corrected using donut correction with a diameter of 9 pixels beyond the detection diameter. In this approach, the mean intensity in an annulus (a donut-shaped region immediately surrounding the particle) is subtracted from the raw particle signal. This compensates for local background fluctuations near each particle and yields the corrected raw mass values used for analysis.

### Particle tracking

Particle coordinates (*x*_1_, *y*_1_, *t*_1_) were linked into trajectories

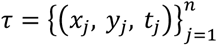

Using TrackPy (46) with a maximum linking distance of 7 pixels and memory of 2 frames. Short trajectories (*n* < *n*_*min*_) with *n*_*min*_ = 14 frames were discarded.

### Distance to membrane (Euclidean distance transform)

For each frame, Euclidean distance transforms (EDT) were computed on both the interior and exterior of the masks. The EDT assigns to each pixel its minimal Euclidean distance to the nearest mask edge. For a particle at (*x, y, t*), the boundary distance was thus

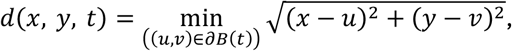

Where *∂B*(*t*) denotes the cell boundary at time *t*. Particles were also classified as inside or outside cells by evaluating *B*(*x, y, t*).

### Summary features and interpretation

From each trajectory we derived descriptors including duration, displacement autocorrelation, fraction of time spent inside cells, mean distance to the boundary, and mean particle intensities. Spatial calibration was applied using the pixel size α (μm/px), and temporal calibration by the acquisition interval Δ*t*.

### Statistical analysis of particle counts, fractions, and sensitivities

All statistical tests were performed on per-biological replicate summary values, rather than individual trajectories, to avoid pseudo-replication. For each replicate and condition, we quantified the number of trajectories at each position (surface, internal, crossing) and derived fractions such as the internalization fraction

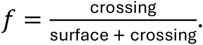

Treatment “sensitivities” were computed per replicate as within-replicate contrasts relative to the IMEM control expressed as differences. Sensitivity was calculated as

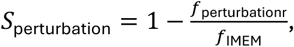

with perturbation being either Pitstop 2 or aptamer here. Pairwise comparisons between two conditions (e.g., variant contrasts within a treatment, or treatment contrasts within a variant) were assessed using a two-sided Wilcoxon signed-rank test applied to the within-replicate differences as it does not assume normally distributed differences. When the Wilcoxon test could not be evaluated due to ties or zero differences, a paired t-test was used as a fallback. p-values are reported directly above bars in figures. This approach controls for replicate-to-replicate baseline variation, provides robustness to non-normal distributions, and ensures that significance reflects biologically interpretable contrasts in particle allocation among entry states.

### Log transform and bootstrap

To test whether two groups of mean step-length values arose from the same underlying distribution, we applied a likelihood-ratio test (LRT) under the log-normal assumption. Each sample was log-transformed and fitted either by a shared normal distribution (null hypothesis) or by separate normals for each group (alternative). The test statistic was the log-likelihood difference. Because the asymptotic reference distribution can be unreliable and overly sensitive for large n, we instead estimated the null distribution of this statistic via a parametric bootstrap: new replicate datasets were simulated from the fitted shared log-normal, and the LRT was recomputed for each. To further reduce spurious significance when sample sizes were very large, both the observed test and bootstrap replicates were calculated on balanced random subsamples capped at 200 trajectories per group. The resulting bootstrap p-value flags only substantial differences in the fitted log-normal parameters: small p (for example, p < 0.05) indicates the groups are unlikely to share a single distribution, whereas large p indicates the data are compatible with a shared distribution at this sample size, not proof of equality (47,48).

## Data and code availability

Data and code will be made fully available upon publication on ERDA.ku.dk and http://github.com/hatzakislab/sarscov2.

## Author contributions

N.S.H. and J.K. conceptualized the project. D.K. produced and characterized VLPs. J.L. and L.C. produced and characterized aptamers. M.G.M. performed NTA measurements. F.H.S. conducted live cell experiments. F.H.S and M.W.D. performed data analysis. F.H.S. wrote the manuscript. All authors reviewed the manuscript. N.S.H. supervised the project.

## Acknowledgements

We thank the members of our laboratories for helpful discussions and encouragement. We thank Emily W. Sørensen for assistance on single particle tracking. This work was funded by the Villum Foundation by being part of BioNEC (grant 18333), The Novo Nordisk foundation challenge center for Optimized Oligo Escape and Control of Disease (NNF23OC0081287), and The Center for 4D Cellular Dynamics (NNF22OC0075851).

