## supplementary info for "Single-Particle Tracking and Positional Phenotyping Reveals Variant-Specific Early Checkpoints in SARS-CoV-2 Cell Entry"

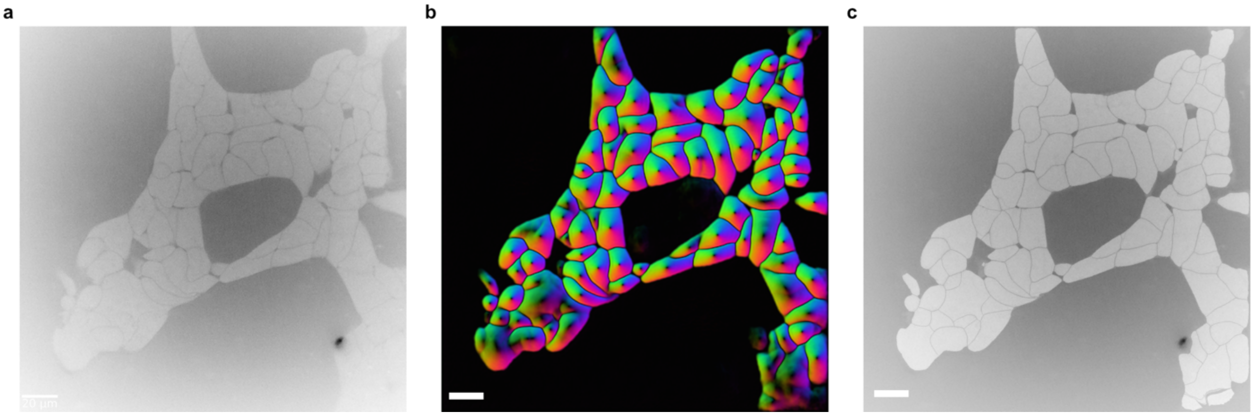


**Supplementary Fig 1. Cell segmentation. a**, Contrast between cells (light grey) and medium (dark grey) was made by addition of 1 μm ATTO647-carboxy to the cell medium. Each cell image was corrected for its illumination profile in preparation for segmentation. **b**, Cell segmentation was performed using Cellpose with a custom model trained a subset of images from the experimental data. Figure shows individually masked cells. **c**, Overlay of cell masks (light grey) overlayed the cell image from **a**. Scale bars, 20 μm.

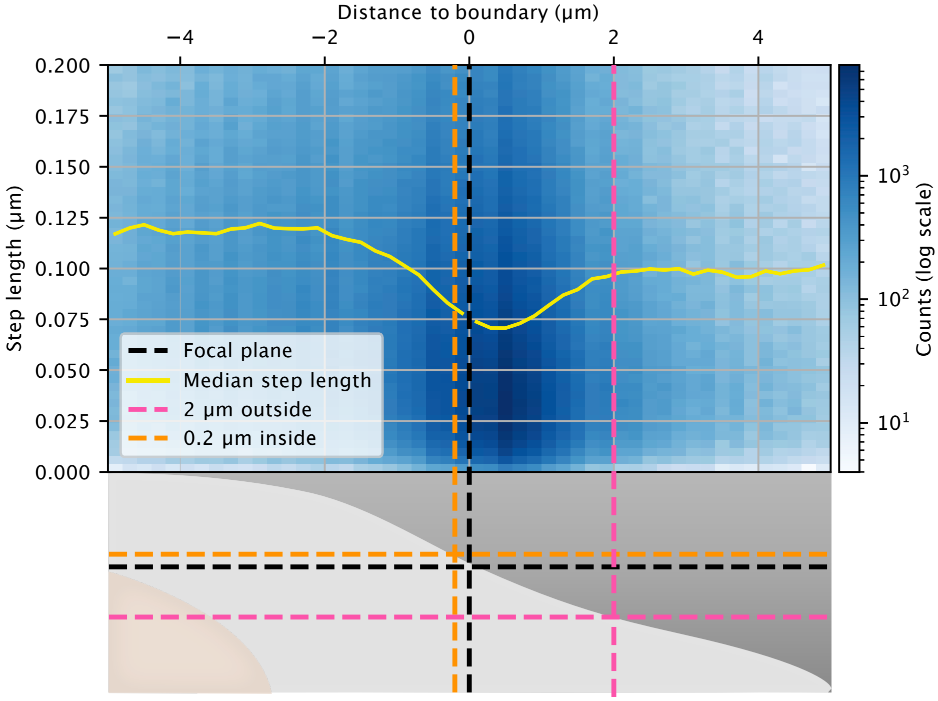


**Supplementary Fig. 2. Determination of distance thresholds for VLP positional phenotyping.** A 2D histogram (blue, log scale) shows the density of VLP trajectory step lengths as a function of the distance to the cell membrane at the microscope focal plane (black lines). The inner and outer distance thresholds (orange lines) that define the outside (membrane-bound) and internal (cytosolic) snapshot positional phenotypes were chosen based on the median VLP step length (yellow line). Step lengths are shortest just outside the center of the cell membrane, where most membrane-bound particles reside. The outer threshold is placed at the distance where the increase in step length reaches a plateau. Using a relatively wide outer threshold allows inclusion of VLPs sitting on flat membrane regions and filopodia that might otherwise be missed during image segmentation. Most unbound VLPs are excluded from the analysis by imposing a minimum trajectory duration of 15 steps, however, some longer trajectories persist as a result of stochastic false linking of fast diffusing VLPs. A small internal distance threshold (magenta lines) ensures that VLPs classified as internal remain internalized throughout their tracked lifetime. Data from all cells were pooled for this analysis.

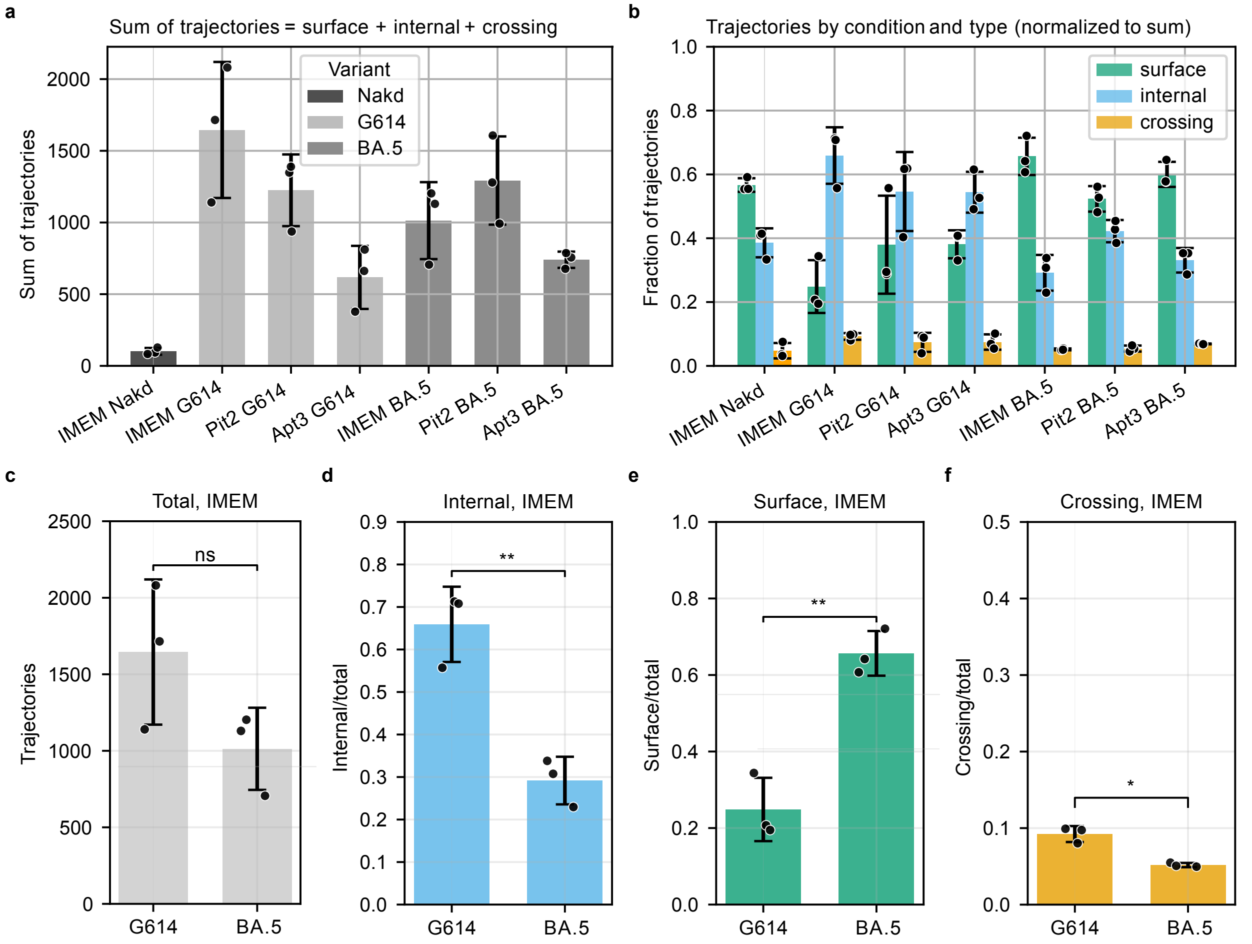


**Supplementary Fig. 3. Total VLP counts. a**, VLP counts were summed for all three positions: surface, internal, and crossing for all conditions. **b**, VLP at all three positional phenotypes for all conditions with each replicate normalized to the sum of the condition. **a-b**, $N_{G614,IMEM}$ = 4936, $N_{G614,Pit2}$ = 3674, $N_{G614,Apt3}$ = 1851, $N_{BA.5,IMEM}$ = 3039, $N_{BA.5,Pit2}$ = 3878, $N_{BA.5,Ap3}$ = 2220. **c,** Total trajectory counts for G614 and BA.5 in IMEM. $N_{G614,IMEM}$ = 4936, $N_{BA.5,IMEM}$ = 3039. **d**, Fraction of internal trajectory counts to total trajectories for G614 and BA.5 in IMEM. $N_{G614}$ = 3332, $N_{BA.5}$ = 914. **e,** Fraction of surface trajectory counts to total trajectories for G614 and BA.5 in IMEM. $N_{G614,IMEM}$ = 1157, $N_{BA.5,IMEM}$ = 1967. **f,** Fraction of crossing trajectory counts to total trajectories for G614 and BA.5 in IMEM. $N_{G614,IMEM}$ = 447, $N_{BA.5,IMEM}$ = 158. Bars show mean ± SEM of three biological replicates. Significance, ns = p>0.05, * = p<0.05, ** = p<0.01.

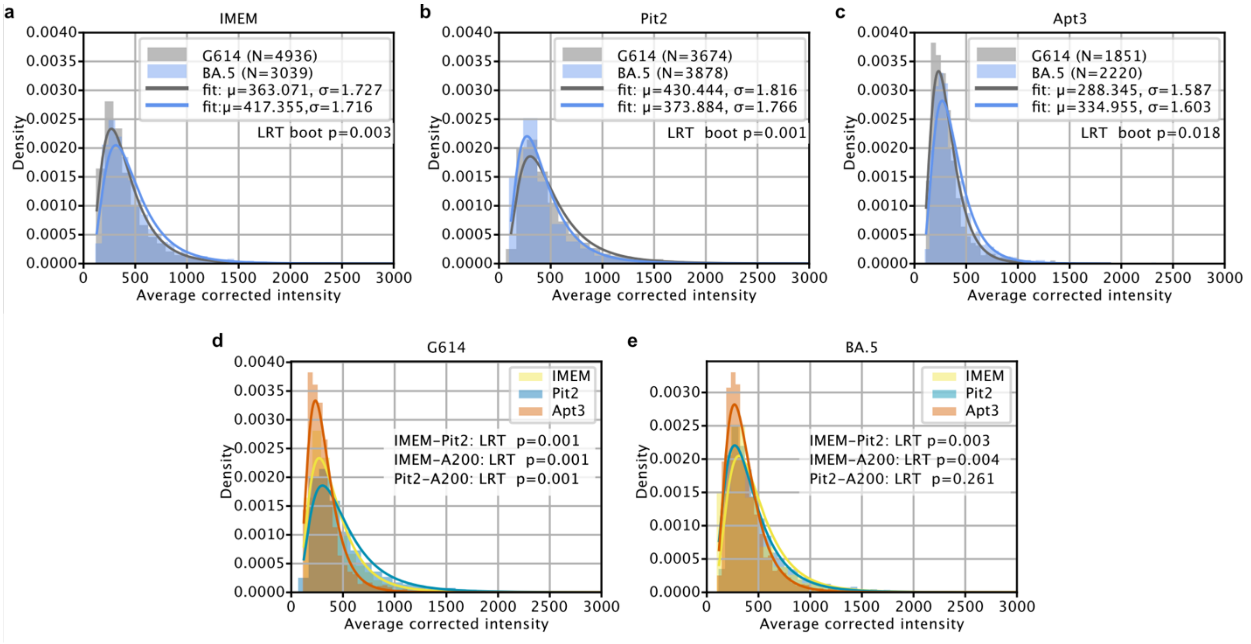


**Supplementary Fig. 4. VLP intensities. a-c**, Comparison of SARS-CoV-2 variant VLP fluorescence intensities between G614 and BA.5 variants under the three conditions IMEM (**a**), Pitstop 2 (Pit2, **b**), and trimeric aptamer (Apt3, **c**). **d-e**, Comparison of single-VLP intensities different experimental conditions of the two spike VLP variants G614 (**d**), and omicron BA.5 (**e**). Distributions were compared using a log-normal likelihood ratio test (LRT) with parametric bootstrap (bootstrap resamples = 1000) and balanced subsampling (subsample size = 200 per group), testing whether two groups share the same log-normal parameters. Reported LRT p-values indicate evidence that both groups share the same underlying log-normal parameters. Data from three biological replicates were grouped per condition.

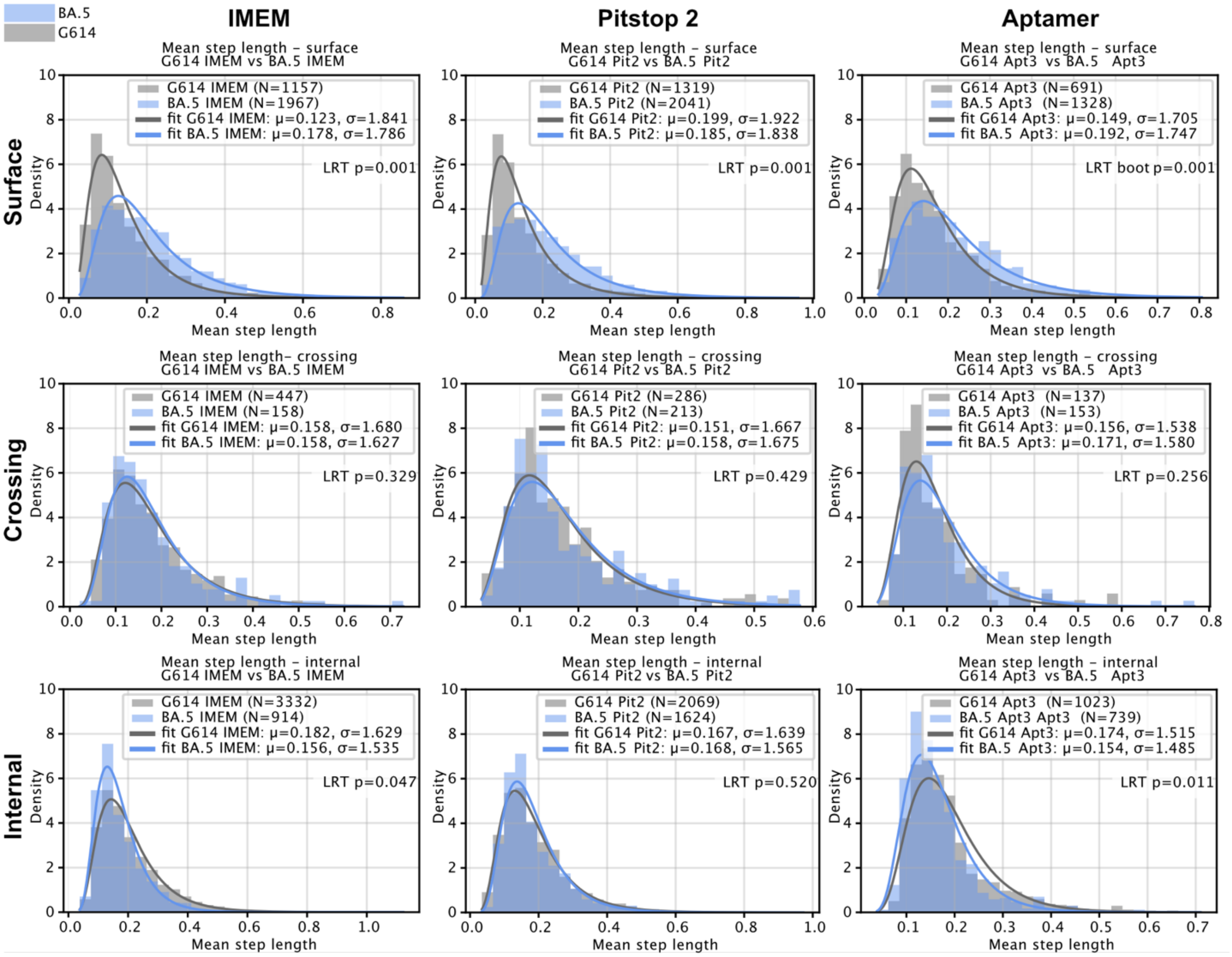


**Supplementary Fig. 5. VLP step length distributions.** Comparison of mean step length distributions between SARS-CoV-2 variant VLPs under different entry conditions and positions. Histograms show per-trajectory mean step lengths with fitted log-normal curves. Variants (G614 vs BA.5) are plotted for each condition: IMEM, Pitstop 2, and trimeric aptamer; and the positional phenotypes surface, internal, and crossing. Distributions were compared using a log-normal likelihood ratio test with parametric bootstrap (B = 1000 resamples) and balanced subsampling (k = 200 trajectories per group). Distributions were compared using a log-normal likelihood ratio test with parametric bootstrap (bootstrap resamples = 1000) and balanced subsampling (subsample size, k = 200 per group), testing whether two groups share the same log-normal parameters. Reported LRT p-values indicate the evidence that both groups share the same underlying log-normal parameters. Curves show log-normal fits to each condition. For each fit, we report the median (exp(μ)) and multiplicative spread (exp(σ)), i.e. the geometric standard deviation). Data from three biological replicates were grouped per condition.
